# The wide distribution and horizontal transfers of beta satellite DNA in eukaryotes

**DOI:** 10.1101/772921

**Authors:** Jiawen Yang, Bin Yuan, Yu Wu, Meiyu Li, Jian Li, Donglin Xu, Zeng-hong Gao, Guangwei Ma, Yiting Zhou, Yachao Zuo, Jin Wang, Yabin Guo

**Affiliations:** Guangdong Provincial Key Laboratory of Malignant Tumor Epigenetics and Gene Regulation, Medical Research Center, Sun Yat-sen Memorial Hospital, Sun Yat-sen University, Guangzhou, China; Institute of Plant Protection and Soil Fertilizer, Hubei Academy of Agricultural Sciences Wuhan China; Department of Parasitology, Zhongshan School Of Medicine, Sun Yat-sen university; Key Laboratory of Tropical Disease Control, Sun Yat-Sen University; Ministry of Education Experimental Teaching Center, Zhongshan School Of Medicine, Sun Yat-sen University; Guangzhou Academy of Agricultural Sciences, Guangzhou, China

**Keywords:** Beta satellite DNA, Sau3A sequences, Horizontal gene transfer, Primates, Eukaryotes

## Abstract

Beta satellite DNA (satDNA), also known as Sau3A sequences, are repeated DNA sequences reported in human and primate genomes. It is previously thought that beta satDNAs originated in old world monkeys and bursted in great apes. In this study, we searched 7,821 genome assemblies of 3,767 eukaryotic species and found that beta satDNAs are widely distributed across eukaryotes. The four major branches of eukaryotes, animals, fungi, plants and Harosa/SAR, all have multiple clades containing beta satDNAs. These results were also confirmed by searching whole genome sequencing data (SRA) and PCR assay. Beta satDNA sequences were found in all the primate clades, as well as in Dermoptera and Scandentia, indicating that the beta satDNAs in primates might originate in the common ancestor of Primatomorpha or Euarchonta. In contrast, the widely patchy distribution of beta satDNAs across eukaryotes presents a typical scenario of multiple horizontal transfers.

**One-sentence summary:** Beta satDNAs in Opimoda could be result of HT from Diaphoretickes and those in primates might have originated in common ancestor of Primatomorpha.

## Main text

The genomes of eukaryotes comprise large tracts of repeated sequences, including satellite DNAs (satDNAs), minisatellite, microsatellite sequences and transposable elements (Charlesworth, Sniegowski, Stephan 1994). Highly homogenized arrays of tandem repeats, known as satDNAs, are enriched in centromeric, pericentromeric, subtelomeric regions and interstitial positions (Yunis, Yasmineh 1971; Greig, Willard 1992). SatDNAs have recently been reconsidered to have various functions, such as playing roles in genome stability, chromosome segregation (Khost, Eickbush, Larracuente 2017) and even gene regulations (Tomilin 2008). Beta satDNAs were considered to be unique in the primates. The basic units of beta satDNAs are 68 bp long with a higher GC content. Beta satDNAs are also known as Sau3A sequences for the presence of a Sau3A restriction site within nearly every single unit (Meneveri et al. 1985). Beta satDNAs have been identified in the genome of human (Meneveri et al. 1993; Eichler et al. 1998), as well as great apes, lesser apes (Meneveri et al. 1995) and old world monkeys (Cardone et al. 2004). However, most of these studies were performed before next generation sequencing was utilized and the amount of sequences studied was quite limited. In this study, we searched genome assemblies of >3,000 eukaryotic species, and revealed that beta satDNAs are widely distributed in eukaryotes. The distribution landscape of beta satDNAs presents a scene of multiple horizontal transfers (HT/HGT) during evolution.

To study the distribution of beta satDNAs in different species, we BLASTed human beta satDNA sequences online against the nucleotide collection of NCBI. In addition to primates that known to harbor beta satDNAs, significant hits were found in *Spirometra erinaceieuropaei, Onchocerca flexuosa, Enterobius vermicularis, Bos mutus* and *Nicotiana tabacum. S. erinaceieuropaei, O. flexuosa* and *E. vermicularis* are endoparasites of human and other mammals. The presence of beta satDNA in these species indicates HT. To our great surprise, hit was also found in a plant, *N. tabacum*. This result suggests that the distribution of beta satDNAs could be much wider than what we have known. It is necessary to perform a global investigation for beta satDNA sequences in eukaryotes.

Currently, there are >8,000 genome assemblies of ∼ 4,000 eukaryotic species in the NCBI Genome database. We downloaded 7,821 assemblies of 3,767 species and BLASTed them against the beta satDNA database containing human beta satDNA sequences. After filtering the BLAST outputs, we found 33,150 beta satDNA copies in 166 genome assemblies of 116 species (Fig. 1, Fig. S1, Table S1 and S2, and Supplementary File 1). Since beta satDNA sequences are highly repeated and difficult to assemble into chromosome contigs, it is hard to assess the ratio of beta satDNAs in certain genomes, e.g. a human genome assembly, GCF_000002125.1_HuRef, has 7,036 beta satDNA copies, whereas, another human genome assembly, GCF_000306695.2_CHM1_1.1, has only 107 copies. In general, beta satDNAs were found in most of the major branches of eukaryotes, including 68 out of 1,394 animals, 16/415 plants, 22/1,656 fungi, 8/60 species of Apicomplexa (9/176 species of Harosa), and 1/20 species of Mycetozoa (1/43 species of Amoebozoa), but no hit was found in Excavata (0/54), which actually is a polyphyletic group (Cavalier-Smith, Chao, Lewis 2015; Brown et al. 2018).

**Fig. 1.**
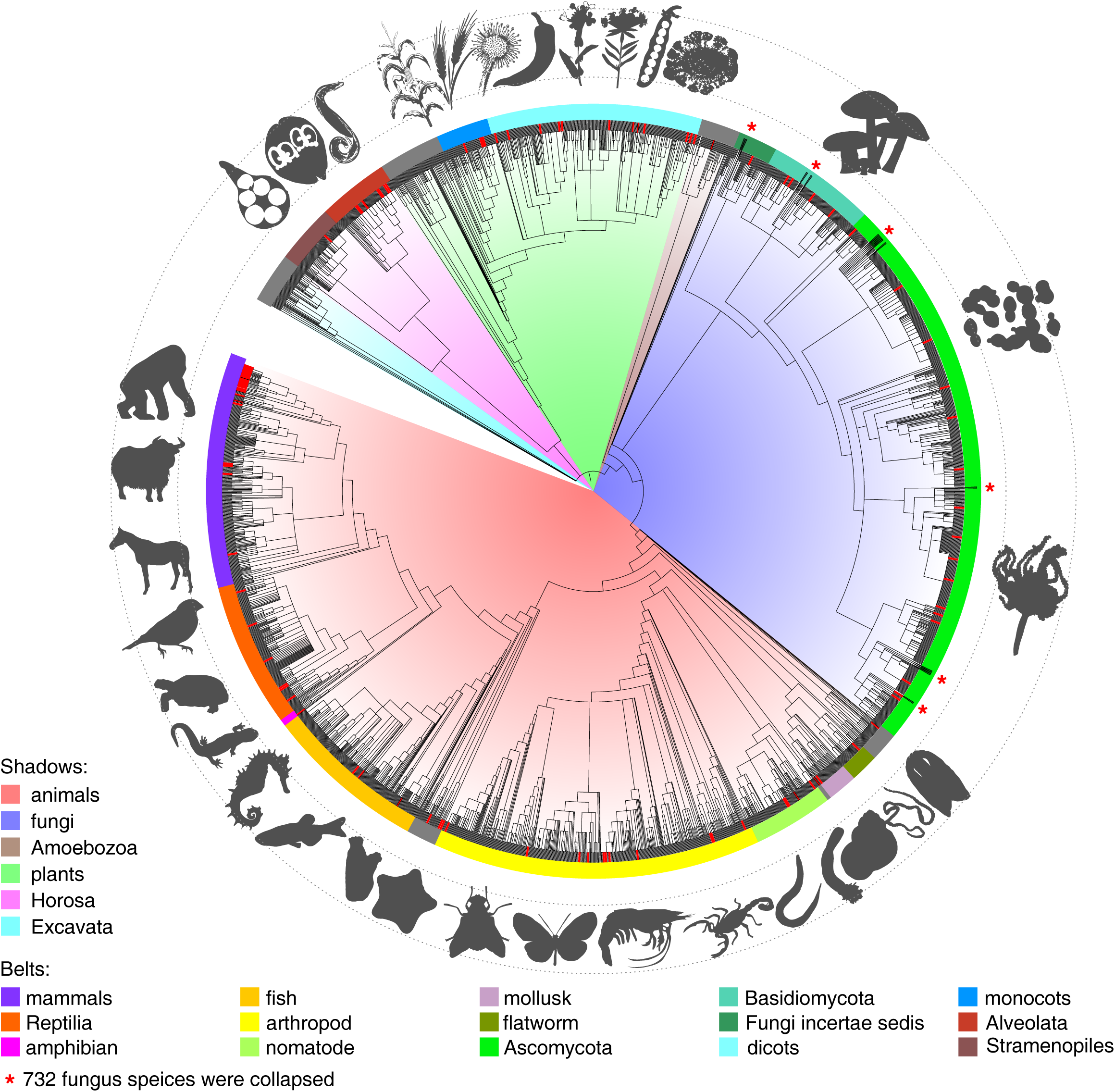
The distribution landscape of beta satDNAs in eukaryotes. 7,821 genome assemblies of 3,767 eukaryotic species were downloaded from the NCBI Genome database and BLASTed against the index of beta satDNA. 116 species (red lines) were found containing beta satDNA sequences.

Beta satDNA can be found in all the major clades of primate, as well as in tree shrew (*Tupaia chinensis*), presenting a vertical transfer scenario. In addition to primates, beta satDNAs were also found in the genome assemblies of four Bovinae species. HT in Bovinae had been reports in several previous studies. It seems that the genomes of Bevinae are especially prone to HT (Kordis, Gubensek 1995; Walsh et al. 2013). Beta satDNAs are rare in the rest of mammals, as well as in other vertebrates and invertebrates (3%), though found in most major taxa. 1.3% fungal species have been found containing beta satDNAs. Since the fungal genomes are usually small and easy to assemble, the existence of beta satDNA in fungi might be significantly rarer than in animals. Interestingly, the proportion of genomes containing beta satDNAs is relatively higher in plants (∼ 3.9%), and even higher in Harosa (5.1%). Moreover, the number of sequences found in Harosa is significantly higher than those in non-primate animals, fungi and plants (Table S1 and Fig. S1). Beta satDNAs typically exist as tandem repeats in human genome, and the similar pattern were also identified in the contigs/scaffolds of certain non-primate species (Fig. S2), indicating that they may play similar roles as they do in primate genomes. Considering that most of the current genome assemblies are far from complete, the actual distribution of beta satDNAs should be substantially wider than the current view, especially in plants, for the extra difficulty in their genome assembling (Claros et al. 2012).

We then analyzed 102 WGS data of 73 species from the NCBI SRA database (Fig.2 and Table S3), so that we can 1) assess the proportion of beta satDNAs in certain genomes; 2) obtain a more elaborate distribution landscape of beta satDNA by looking at some representative nodes on the tree of eukaryotes. Similar to the genome BLAST results, beta satDNAs were found in all primate clades, as well as in Dermoptera and Scandentia. Previous study suggested that there are two bursts of beta satDNA through the evolution of Hominidae: one is after the separation between great apes and lesser apes, and the other is after the separation between African great apes and orangutan (Cardone et al. 2004). However, our result showed that the proportion of beta satDNAs in orangutans is similar to those in gibbons or old world monkeys. Thus, the ‘big bang’ of beta satDNAs was taken place in the African apes, which may suggest a distinct aspect of heterochromatin regulation in the Homininae. Besides Euarchonta, bovines, toothed whales and elephants were the three mammalian clades found having beta satDNAs. Beyond mammals, beta satDNAs were identified in many parasites including blood-feeding insects. Since it is always hard to avoid human DNA contamination in isolating parasites DNA samples (Oyola et al. 2013), this observation need to be treated with special caution. The existence of beta satDNAs seems more common in plants than in animals, though the abundances are not high. We found significant hits in most major taxa of plants, including Angiosperm, Gymnosperm, moss, green algae and red algae. Additionally, the three species of Harosa checked here all contain beta satDNA sequences. Generally speaking, the current WGS data of protists are not many and the quality is not high. We expect a better prospect of beta satDNA in protists in the future.

**Fig. 2.**
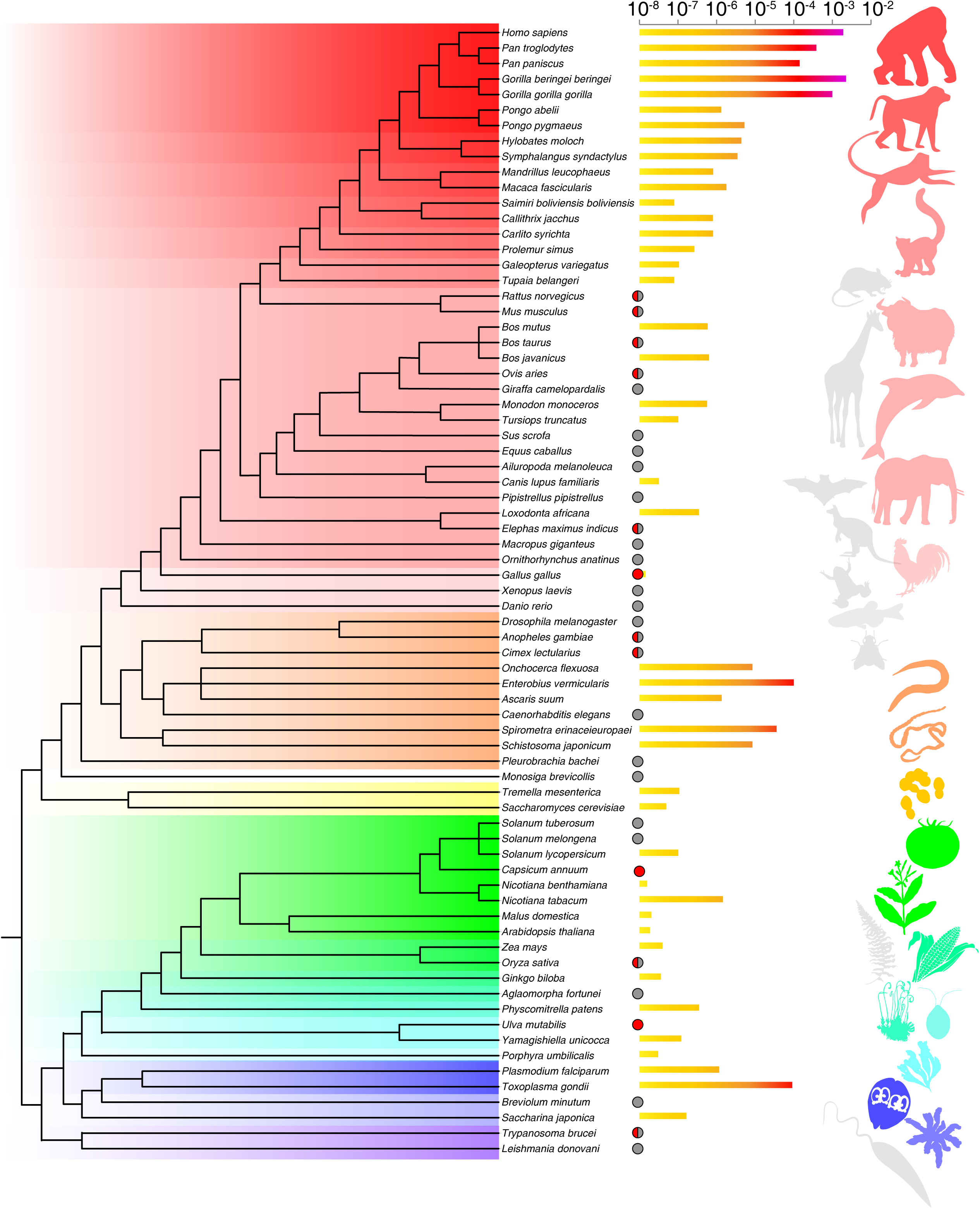
Beta satDNAs in WGS data (SRA). 102 SRA datasets of 73 species were BLASTed against the beta satDNA index. The proportions of beta satDNAs in different species were shown by the histogram at the right part. The evolutionary relationships of the species were shown at the left part. Color range: red, vertebrates (from light to dark: vertebrates, mammals, Euarchonta, primates, Simiiformes, Catarrhini, apes and Hominidae); orange, invertebrates; cyan-green, plants (from cyan to green: plants, green plants, Embryophytes, Equisetophyta, Equisetophytina, Spermatophytes, Angiosperms, dicots); blue, Harosa (dark, Alveolata); purple, Euglenozoa. Circles: gray, not found or ratio <10^−8^; red, found but the bars are too short to view; half red half gray, found in certain SRAs but not found in others.

To validate the search results in genome assemblies and SRAs, we performed PCR assays using genomic DNAs (gDNAs) of 36 species as templates (Fig. 3 and Fig. S3). The positive PCR signals of beta satDNAs are ladders with ∼ 70-bp spacing for their tandem repeated pattern. Basically, the PCR results were consistent with the BLAST results. The signals of mouse, rat and pig showed negative as the negative controls, while a wide range of positive signals were observed in various animal and plant genomes.

**Fig. 3.**
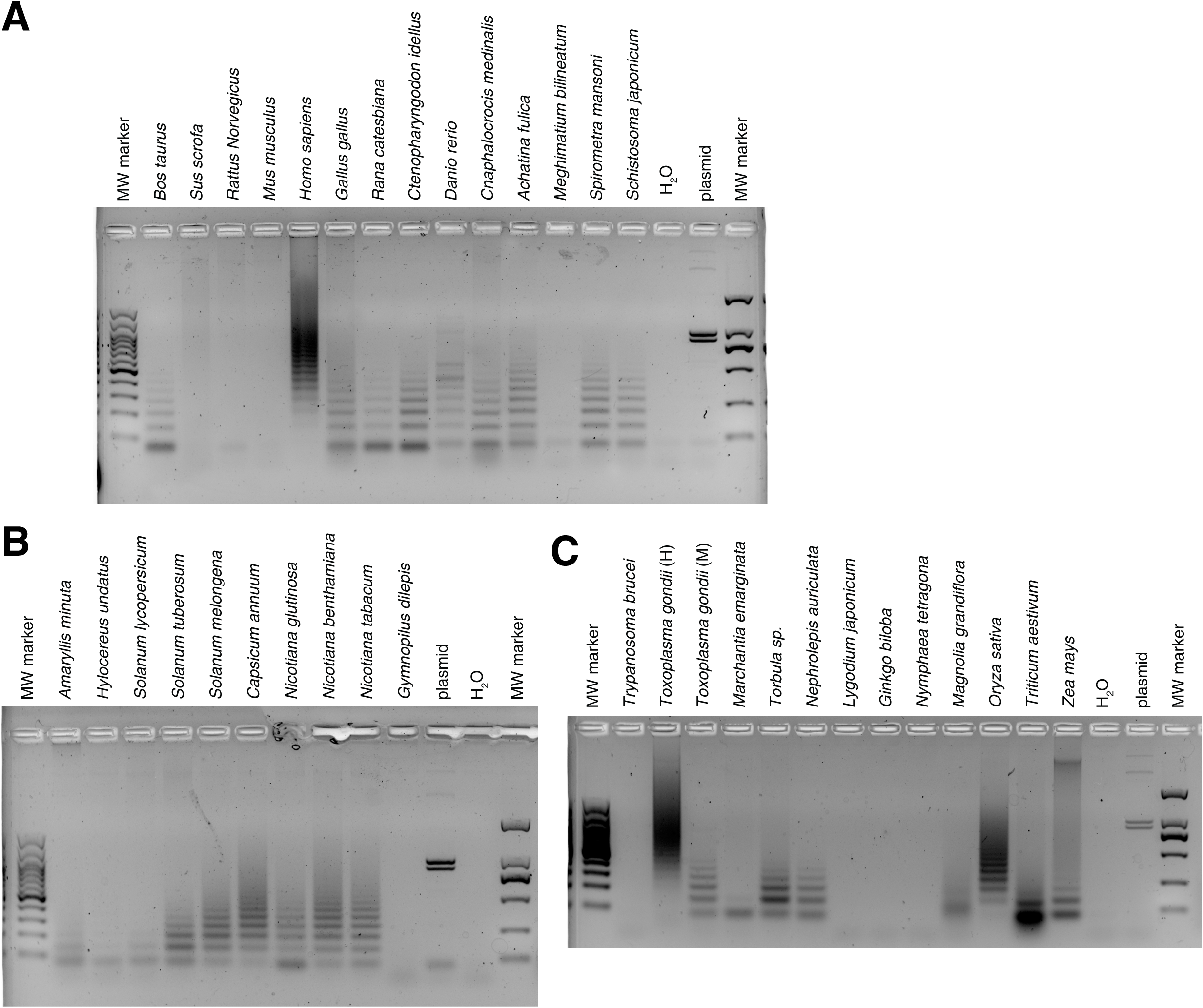
PCR amplifications of beta satDNAs. Genomic DNA samples were isolated from certain organisms and used as templates. Primers were designed according to the consensus sequence of beta satDNA. The products were then analyzed using electrophoresis with 1.5% agarose gel containing gel-red.

It is not surprising that the positive ratio of beta satDNA in SRA, and especially in PCR are significantly higher than that in the genome assemblies. Since beta satDNA sequences are difficult to assemble for their high repetition, many assemblies don’t contain beta satDNA sequences, though the physical genomes do, e.g. we even failed to identify beta satDNA sequences from the current genome assemblies of *Pongo pygmaeus* (Bornean orangutan), who no doubt has beta satDNAs.

There are concerns of contamination in both WGS and PCR, and even the genome data have possibility of contamination and wrong assembling. Therefore, we tried multiple approaches to rule out the risk of false conclusion introduced by possible contamination (see Supplementary Text for detailed information).

The phylogenetic relationships of the beta satDNA sequences from different species were also examined (Fig. 4A). Unlike the case in coding genes or transposable elements (TEs), the beta satDNA sequences from different species couldn’t be separated in phylogenetic tree, which may be due to the intrachromosomal and interchromosomal exchanges (Rudd, Wray, Willard 2006). For this reason, the HT pathways of beta satDNAs cannot be determined like those of coding sequences based on the phylogenetic relationship of sequences from different species. Then we compared the sequence libraries of different species pairwisely (Fig. 4B). Clearly, the diversity of sequences in primates is higher than the rest, indicating that the pool of beta satDNA is quite small until it bursted in primates. Similarly, the cluster analysis on the beta satDNA sequences of non-primate species showed distinct pattern from that on the total sequences (Fig. S4 and S5).

**Fig. 4.**
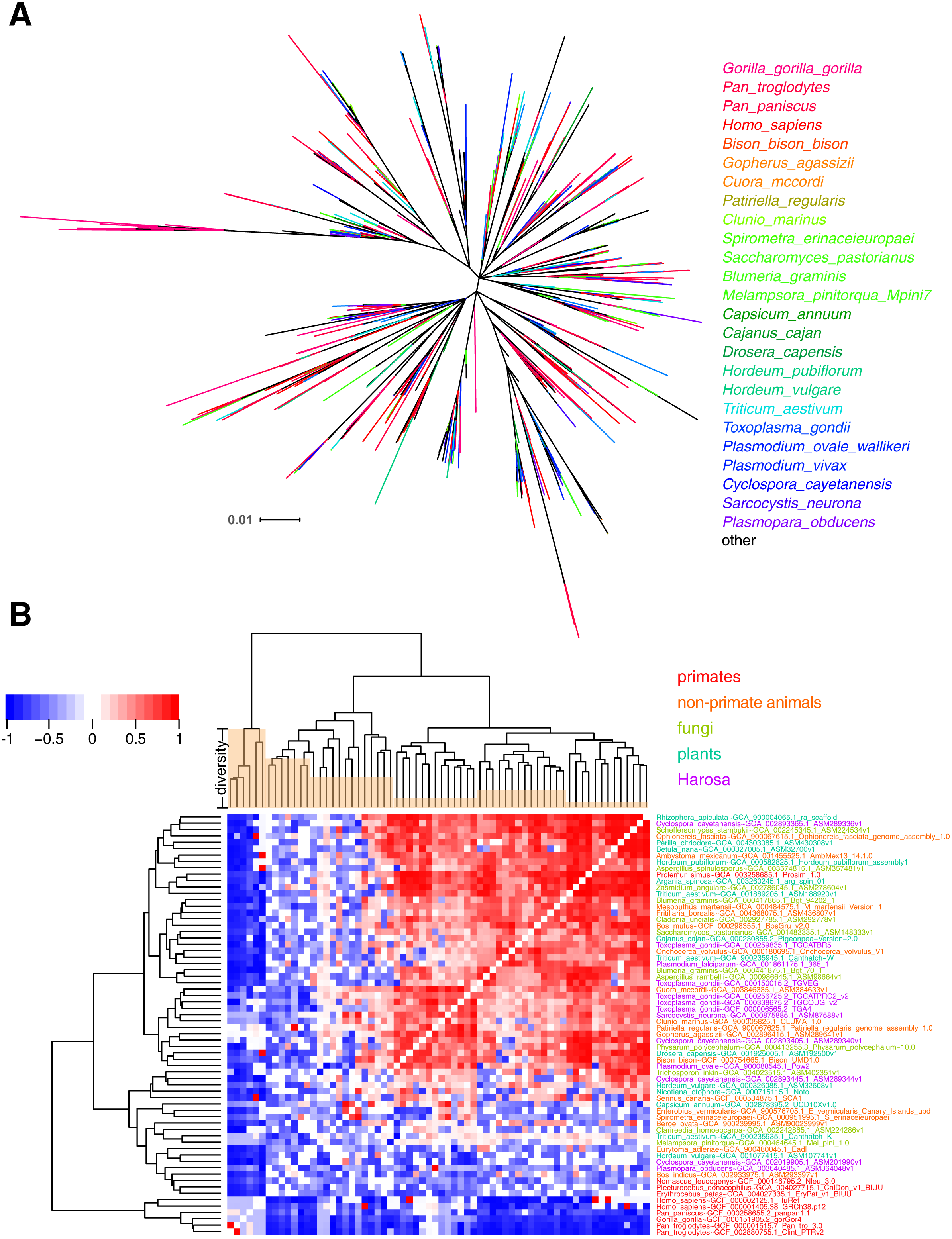
The relationship between beta satDNA sequences identified from genome assemblies of different species. a, Phylogenetic tree of 1,384 beta satDNA sequences of 92 species was constructed using RAxML, with statistical support provided by bootstrapping over 1,000 replicates. b, The sequence libraries of different species that contain ≥ 10 beta satDNA copies were compared pairwisely. The matrix was normalized by a matched random control, and then heatmap was produced using heatmap.2 in R language.

We also searched the 210,000 prokaryotic genome assemblies (12,398 species of 569 archaea and 11,829 bacteria) in the NCBI Genome database and found beta satDNAs in 72 species, two archaea and 70 bacteria (Fig. S6). Obviously, beta satDNAs in prokaryotes are far rarer than in eukaryotes. Most of these bacteria are symbionts/pathogens of animals or plants (Table S4) and they might acquire beta satDNAs from their hosts. In addition, there are >13,000 genome assemblies of organelles and >30,000 genome assemblies of viruses in the NCBI database, but none of them contains beta satDNA. Although viruses and symbiotic organelles are common media for HT (Bergthorsson et al. 2003; Liu et al. 2010), they might not play roles in the HT of beta satDNAs.

Employing multiple approaches, we obtained a high-resolution distribution landscape of beta satDNAs in eukaryotes. It is previously thought that beta satDNAs were originated in Catarrhini and bursted in great apes (Cardone et al. 2004), whereas, our study showed that beta satDNA actually exists widely across eukaryotes. The patchy distribution of beta satDNAs in animals and fungi suggests multiple HT events during evolution. However, the existences of beta satDNA seem wider in plants, and especially in Harosa (Table S1 and Fig. S1). Animals, fungi and Amoebozoa belong to Opimoda, while plants and Harosa belong to Diaphoretickes. All the beta satDNA sequences identified here are within Opimoda and Diaphoretickes (Fig. 1 and 2). No full-length beta satDNAs were found in any branches of Excavata so far, with only several truncated fragments found in *T. brucei* (Euglenozoa). Therefore, we hypothesize that beta satDNA was an ancient sequence that originated in Diaphoretickes, after its separation with Euglenozoa. The beta satDNAs in Opimoda were diffused from Diaphoretickes via HT. Parasites play critical roles in HT as reported previously (Deitsch, Driskill, Wellems 2001; Gilbert et al. 2010). The members of Apicomplexa are all kinds of parasites of animals, including the pathogens of malaria, toxoplasmosis, cyclospriasis etc., and they might have played the major role of transferring beta satDNAs to animals. Of course, today’s parasites of Apicomplexa have been co-evolving with their hosts through all the time, so that the current contents of beta satDNAs in their genomes should be the results of countless two-way HT events. The distribution of beta satDNA in Plants could be the common result of inheritance from ancestors, loss and HT. Most of the fungi that were found to have beta satDNAs are parasitic and a few saprophytic. They might have acquired beta satDNA from their hosts of the plants or animals in their living environments.

We identified beta satDNA sequences in primates, as well as in Dermoptera and Scandentia, but not in mouse, rat or other rodents. Given that the mouse genome has been thoroughly studied, we tend to believe that the mouse genome doesn’t comprise beta satDNAs and beta satDNAs are absent or at least not common in rodents. Scandentia (tree shrew) was previously considered a sister group of Primatomorpha, but recently was reconsidered a sister group of Glires (Meredith et al. 2011; Zhou et al. 2015). Thus, the origin of beta satDNAs in primates could be traced back to at least the common ancestor of Primatomorpha (∼ 80 MYA).

HT/HGT was considered one of the major drives of evolution, while recent investigations suggest that the role of HT may be even more important than had been thought (Keeling, Palmer 2008; Daubin, Szollosi 2016). Complex eukaryotic genomes usually comprise large fraction of repeated sequences, including TEs and non-transposable elements. Previous studies have shown that HT is important in the origin of TEs in certain genomes (Gilbert et al. 2012; Walsh et al. 2013; Gilbert, Feschotte 2018; Ivancevic et al. 2018), while here we suggest that HT may be important for the origin of satDNAs too. As the deep sequencing capacity expands tremendously, the main stream of the studies on HGT will certainly transit from studies based on special cases to systematic studies (Gilbert et al. 2010; Ivancevic et al. 2016; Peccoud et al. 2017; Ivancevic et al. 2018). Only by this way can we evaluate the great impact of HT for evolution more accurately. We believe a new distribution landscape of beta satDNAs with higher resolution will be obtained in a few years when more genome assemblies with high quality are available.

Our study greatly updated the knowledge on the distribution of beta satDNA in the tree of life and the origin of the beta satDNAs in primates. The previous investigations on HT were mainly focus on certain genes or TEs. Here we showed that satellite DNAs too are materials for HT, and enriched the topics of the HT research field. Furthermore, the beta satDNAs found in some laboratory models, such as certain species of budding yeast, fruit fly and tobacco, suggest that studies on beta satDNA can be performed in these simple organisms in the future, besides in primate cell lines.

## Supporting information

beta satDNA sequences

