## Supplementary text, methods, table and figures for "The wide distribution and horizontal transfers of beta satellite DNA in eukaryotes"

**Supplementary Materials**

Supplementary Text

Materials and methods

Supplementary References

Supplementary Table 1

Supplementary Table 2-4 and 5 (see the Excel files)

Supplementary Figure 1-10

Supplementary File 1 (see the fasta file)

### Supplementary Text

#### Beta satDNAs in human genome

In the human genome, Beta satDNAs have been identified in the pericentromeric regions of chromosomes 1, 9, and Y (Meneveri et al. 1993), as well as in acrocentric chromosomes 13, 14, 15, 21 and 22 (Agresti et al. 1987) and in chromosome 19p12 (!!! INVALID CITATION !!!). Beside the roles in genome stability and chromosome segregation (Khost, Eickbush, Larracuente 2017), beta satDNAs may even involve mammalian gene regulations (Tomilin 2008). The changes in the copy number of repetitive sequences in genomic DNA are important causes of hereditary disorders (Mirkin 2006; Mirkin 2007; J. Rich, V. V. Ogryzko, Pirozhkova 2014). The insertion of 18 beta satellite units in the gene coding a transmembraine serine protease causes congenital and childhood onset autosomal recessive deafness (Scott et al. 2001). The global DNA hypomethylation frequently observed in cancers is mostly taken place at satDNAs (J. Rich, V. V. Ogryzko, Pirozhkova 2014). The loss of the BRCA1 tumor suppressor gene provokes satDNA derepression in breast and ovarian tumors in both mice and humans (Zhu et al. 2011). Facioscapulohumeral muscular dystrophy (FSHD), a autosomal dominant hereditary disease, is associated with the macrosatellite (D4Z4) and beta satellite (4qA allele) DNA sequences (J. Rich, V. V. Ogryzko, Pirozhkova 2014).

The current human genome assembly is not yet complete for the high degree of sequence homogeneity among many hundreds or thousands of copies of repeated sequences. Most human chromosome contigs contain few or no beta satDNA sequences except chromosome Y. The beta satDNA sequences on chromosome Y are clustered in three separated regions designated as Ya, Yb and Yc in this study (Fig. S7A). The sequences of the three regions were mapped and single units of beta satDNA sequences were extracted. After CLUSTALW alignments being performed, the phylogenetic relationships of these sequences were examined using RAxML software (Fig. S7B).

Obviously, the sequences from Ya, Yb and Yc are not phylogenetically separated. To see if the beta satDNA sequences on chromosome Y are different from those on other chromosomes, we randomly picked 1000 copies of non-repeated beta satDNA sequences from the human beta satDNA dataset to build a phylogenetic tree together with the beta satDNA sequences from chromosome Y (Fig. S7C). This result shows that there is no regional or chromosomal specificity in human beta satDNA sequences, indicating that the evolution of beta satDNAs, similar to the higher-order alpha satDNAs, are extremely homogeneous within and between chromosomes, which may be a result of intrachromosomal and interchromosomal exchanges (Rudd, Wray, Willard 2006). The beta satDNA sequences on chromosome Y have relatively higher diversity among the beta satDNA sequences in human genome, which is consistent with the previous report (Cardone et al. 2004).

The unequal exchange is a strong long-range ordering force which can keep tandem arrays homogeneous (Charlesworth, Sniegowski, Stephan 1994). The homogenization process is much faster in the bisexual species, suggesting that meiotic recombination accelerates the homogenization process (Mantovani et al. 1997). Concerted evolution leads to higher homogeneity between the satDNA sequences of intraspecies than the sequences of interspecies (Dover 1982; Kuhn et al. 2007). The higher-order alpha satDNAs are significantly more conservative within species than between primate species (Rudd, Wray, Willard 2006; Cacheux et al. 2016). However, the beta satDNA sequences lack obvious conservative property within species (Fig. 4), indicating the absence of complete concerted evolution, which might be related to a presumed reduction or suppression of meiotic recombination (Kuhn et al. 2007).

#### The possibilities of artifacts in genome assemblies, SRA and PCR

Since we are the first to report beta satDNAs in non-primate species, we treated the data with special caution to rule out the possibilities of artifacts.

Theoretically, the beta satDNA copies found in non-primate genome could be wrong assemblies of sequence reads contaminated by human DNA. Presumably, the contamination would occur randomly. However, the distribution of beta satDNA in life tree is highly aggregated. For example, a) in mammals, beta satDNAs are found almost only in primates and bovines. Four species of Bovidae/Bovinae were found containing beta satDNAs. b) Beta satDNA seems more common in plants than in animals (not counting primates), even though the plant genomes are more difficult to assemble than those of animals(Claros et al. 2012). c) The positive ratio and the total sequence number of beta satDNA in Apicomplexa is significantly high than in other taxa. Many members of Apicomplexa are human parasites and their gDNA samples have the risks of contaminated by human DNA. However, there are many parasites in Excavata/Discoba too, yet none of the genomes in these species were found containing beta satDNAs (Fig. 1). Moreover, the positive ratio for beta satDNAs in fungal pathogens is not high as well. Therefore, the observation of beta satDNAs in Apicomplexa is unlikely to be an artifact.

Furthermore, the beta satDNA sequences found in different genome assemblies of one species or in different species of one genus made the finding more credible. For example, beta satDNAs were found in six genome assemblies of three butterfly species in the *Heliconius* genus, six assemblies of two species in *Hordeum* genus, five assemblies of *Triticum aestivum,* 11 assemblies of *Toxoplasma gondii* and assemblies of four species in *Plasmodium* genus, etc (Table S2).

If the beta satDNAs found in the assemblies of non-primate species were due to contamination of human DNA, there should be no specificity between the sequences found in assemblies of different species. However, Fig. 4B and Fig. S4-S5 showed that these sequences can be grouped into certain clusters. In contrast, when the sequences of different species were randomized and the analysis was performed again, clustering was failed and the order of species was completely different (Fig. S8).

There are concerns of contamination in both whole genome sequencing and PCR. We tried multiple approaches to rule out the risk of contamination. a) The SRA datasets were screened manually to make sure most of the datasets containing human DNA < 0.01% (or satisfying similar criteria). b) The parasite samples used for isolating genomic DNA for PCR were all taken from non-primate animals, but not from humans. c) PCR assay was performed with a series diluted human DNA as template and they always produce large molecular weight ladders even the template concentrations are extremely low (Fig. S9). PCR were performed using gDNAs of *T. gondii* from both cultured human cells and mice as templates, which showed distinct ladder patterns in agarose gel (Fig. 3C, lane 3 and 4). These results indicate that if a sample is contaminated by human gDNA, it will show a typical “human style” ladder, which we didn’t see in the PCR with gDNA templates of other species.

The PCR amplicons of human, cattle, tapeworm, blood fluke, tomato and tobacco were also submitted for next generation sequencing. The sequence libraries of these species were compared pairwisely. Fig S10 showed that the libraries are distinct. Similar to Fig 4B, the beta satDNA sequences of human has the largest diversity, while the libraries of the two plants of Solanaceae showed the highest similarity. This result showed that the sequences were amplified from different source templates, instead of contamination of human gDNA.

Finally, the results of genome BLAST, SRA BLAST and PCR assay were highly consistent. Controversial results were only observed in Zebra fish (negative in SRA BLAST, but positive in PCR) and ginkgo (positive in SRA BLAST, but negative in PCR). Although the possibility of human gDNA contamination always exists, a large fraction of beta satDNA sequences found in non-primate species should be *bona fide*. Therefore, we believe that beta satDNAs do exist across eukaryotes.

**Summary**

In summary, a) we performed the ever largest-scaled study on beta satDNAs. b)

The origin of beta satDNA in primates can be traced back to the common ancestor of Primatomorpha or Euarchonta (at least 80 MYA). c) Beta satDNAs widely exist across eukaryotes. d) The HGT of satellite DNAs was shown for the first time. e) Our study raises new topics for the research fields of satellite DNA and HGT.

### Materials and Methods

#### Identification of beta satDNAs from genome assemblies

All the available genomic sequences (fasta files) of eukaryotes, prokaryotes, organelles and viruses in the National Center for Biotechnology Information (NCBI) Genome Database (https://www.ncbi.nlm.nih.gov/genome/browse#!/overview/) were downloaded (Table S5), except some assemblies that were failed to download or of low quality. The sequences were BLASTed against an index built with pre-identified beta satDNA sequences of human and other apes. The BLAST outputs were filtered using a Perl script, and sequences with match length $\geq$ 55 and e-Value $\leq 10^{-9}$ were kept. Sequences with match length = 69 and TC/GA (Sau3A restriction site) at start/end were designated as full-length beta satDNA sequences.

#### Identification of beta satDNAs from human chromosome Y

The full sequence of the chromosome Y (NC_000024.10) was downloaded from NCBI. The locations of beta satDNAs on chrY were determined by making preliminary comments using Geneious 11 (Kearse et al. 2012), and the three regions containing beta satDNA copies, Ya, Yb and Yc were identified. Then the sequences of the three regions were mapped to a 68 bp beta satDNA reference sequence based on a 72 bp sliding window at 1 bp resolution (e.g. nt 1-72, nt 2-73…) using Geneious 11, and the dataset of the beta satDNA sequences on chrY was obtained. The scripts for extracting sequences were written in Python language.

#### Identification of beta satDNAs in the WGS data of different species

The raw sequencing data of the 73 species were downloaded from the NCBI SRA database (https://www.ncbi.nlm.nih.gov/sra) or BLASTed on line. Then the BLAST output files were filtered with the same criteria as the genomic BLAST outputs, and the beta satDNA sequences were extracted using scripts written in Python. The percentages of beta satDNAs in each species were calculated using the following formula:

```
number of hits / total number of reads in SRA databases
```

#### Generation of trees of life

The trees of life (Fig. 1, Fig. 2, Fig. S3 and Fig. S6) were originally generated at the Common Tree webpage of NCBI Taxonomy (https://www.ncbi.nlm.nih.gov/Taxonomy/CommonTree/wwwcmt.cgi). The .phy tree file were then modified manually and further edited at iTOL (https://itol.embl.de/) (Letunic, Bork 2016).

#### Phylogenetic analyses

Sequence alignment was performed using the MUSCLE(Edgar 2004), All phylogenetic analyses were conducted under maximum likelihood in RAxML (Stamatakis 2014). The phylogenetic trees were colored at iTOL (https://itol.embl.de/).

#### Isolations of genomic DNAs

The larvae of *Spirometra mansoni* (plerocercoid) were collected from frogs. *Trypanosoma brucei* and *Toxoplasma gondii* were collected from cell culture medium or mice. The adult *Schistosoma japonicum* were collected from rabbits. The genomic DNAs of the animals were isolated using TIANamp Genomic DNA Kit (TIANGEN). The genomic DNAs of plants and fungi were isolated using Plant DNA Mini Kit (OMEGA).

#### PCR assay

PCRs (20 μl) comprised 2 μl genomic DNA (10–80 ng), 2 × Premix Taq (TaKaRa), 0.5 μl forward / reverse primers (see below for sequences). The PCR cycling program was 30 cycles of 98 ℃ for 10 s, 56 ℃ for 30 s, 72 ℃ for 1 min and a final extension step of 72 ℃ for 10 min. The PCR products were checked by electrophoresis with 1.5% TAE agarose gel containing gel-red.

Primer F: GATCACCCAGGTGATGTAACTCTTGTC

Primer R: GATCAGTGCAGAGATATGTCACAATGCC

**Next generation sequencing of amplicons**

Barcoded primers were used in PCR for amplifying beta satDNAs from different gDNA samples. The amplicons were purified and sequenced at Guangzhou IGE Biotechnology. The raw sequence reads were then BLASTed against the beta satDNA index and filtered under the same parameter as the genome and SRA BLAST filtering.

**Cluster analysis of beta satDNA sequences**

Full-length beta satDNA sequences were compared pairwisely. The matrix of identities was used for the clustering analysis and heatmap generation using the heatmap.2 function of ‘gplots’ library in R language.

**Cluster analysis of sequences identified from different genome assemblies**

For the genome assemblies containing beta satDNA sequences, those that contain ≥10 beta satDNA copies were chosen. The beta satDNA libraries were compared pairwisely and the common ratios were calculated using the algorithm as below:

R-common = (libA ∩ libB)/(libA ∪ libB)

And a matrix of R-common was generated. Considering the sizes if libraries varies greatly, a matched random control matrix was created by randomizing the sequences and ran the script for 20 times to get an average. Then the difference of the two matrixes was used for the clustering analysis and heatmap generation using the heatmap.2 function in R language. GCF_000306695.2_CHM1_1.1 and GCA_002754635.1_CMB-1_v2 were excluded in Fig. 4 for very different copy numbers in relative genera.

**Table S1.** A summary of beta satDNA sequences found in the genome assemblies of eukaryotes.

| Taxa | Species | copy number | copy number of full-length units |
|---|---|---|---|
| primate | 28 | 29067 | 16140 |
| animal excluding primate | 40 | 673 | 445 |
| fungus | 22 | 317 | 252 |
| Amoebozoa | 1 | 19 | 17 |
| plant | 16 | 435 | 321 |
| Harosa | 9 | 2639 | 2103 |
| total | 116 | 33150 | 19278 |
| unique seqences | | 18390 | 10966 |

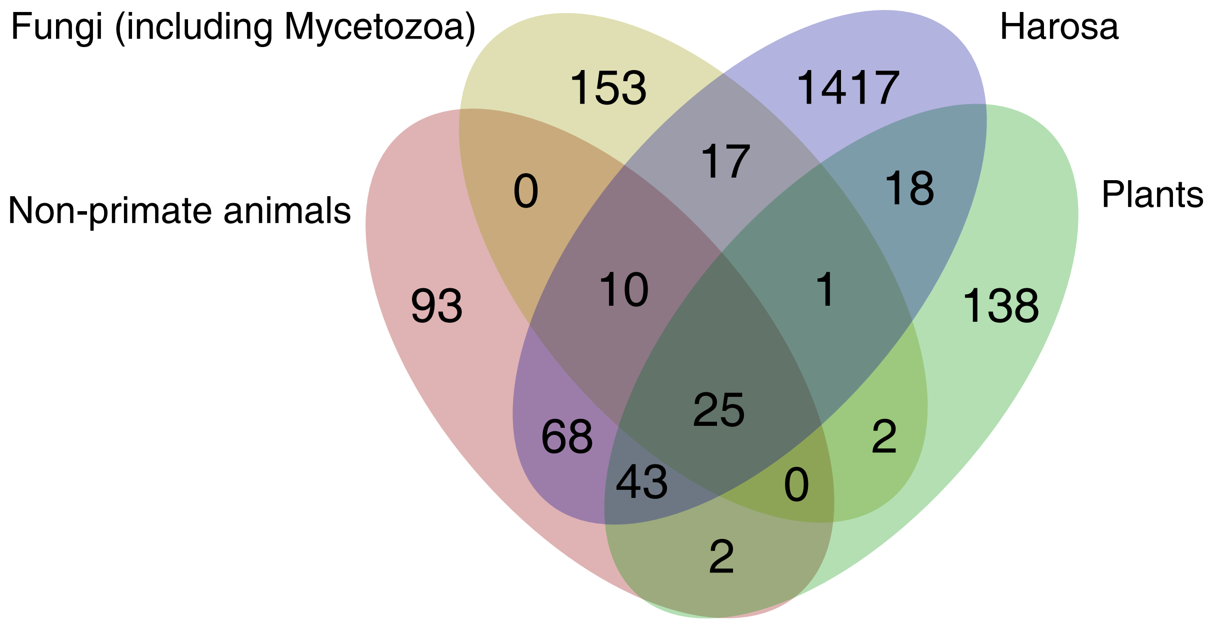


**Fig. S1.** Venn diagram shows the number of full-length beta satDNA sequences found in four major eukaryotic branches.

**Fig. S2.** Tandem repeats of beta satDNA units identified in contigs of genome assemblies. The chromosome contigs of certain species were randomly selected, and the beta satDNA units (brown triangles) on the contigs were labeled using Geneious 11. The species and the contig Ids were marked on the left.

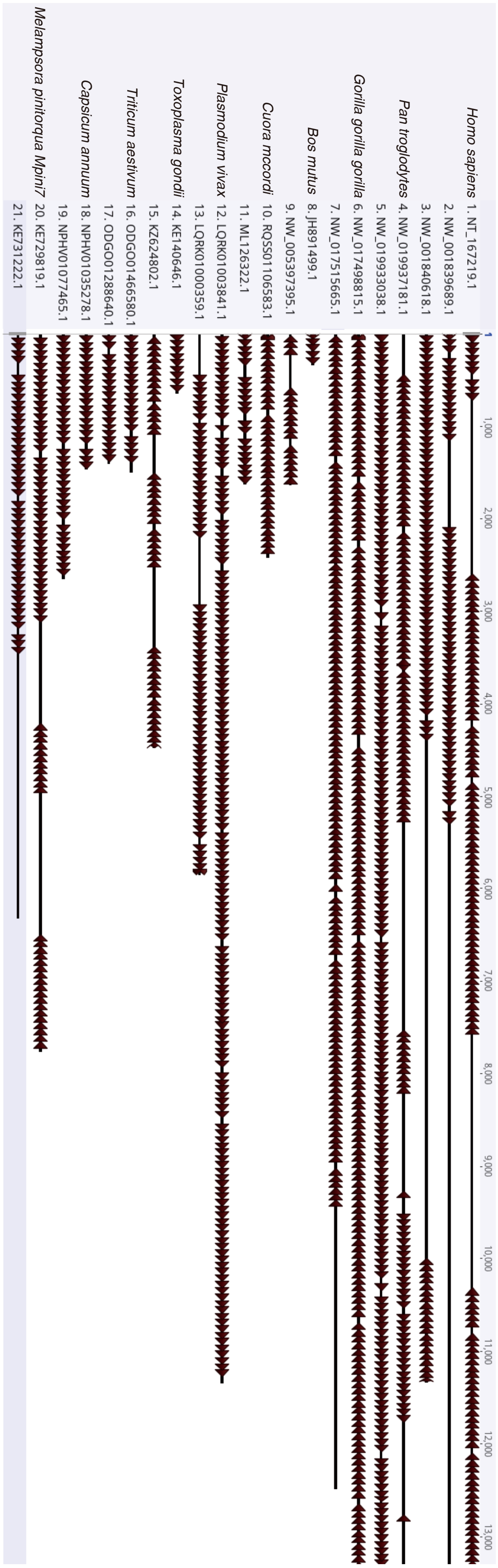

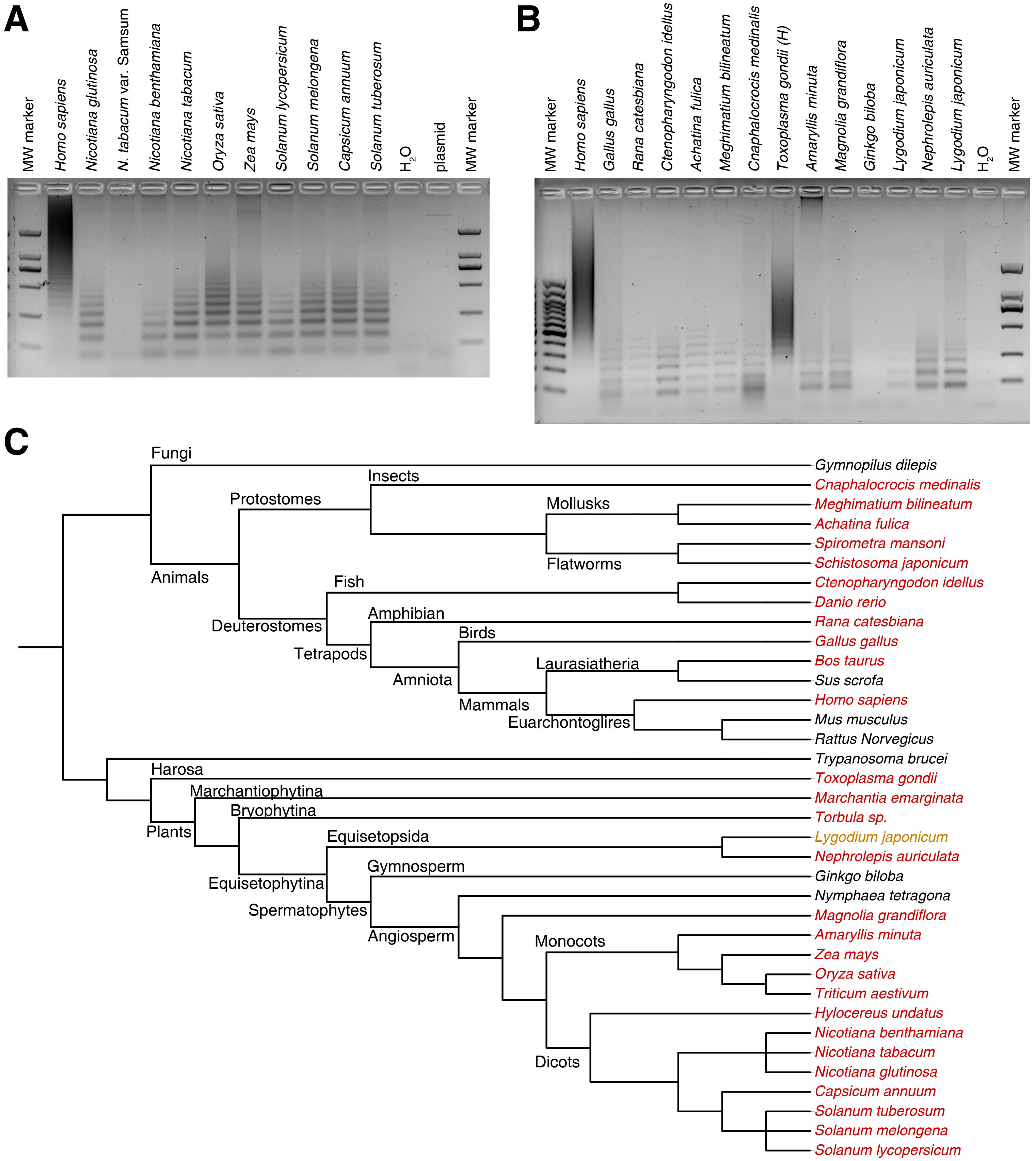


**Fig. S3.** PCR amplifications of beta satDNAs. A and B, An extension of Fig. 3. C, The evolution relationships of the 36 species tested in PCR assays. Red, positive; black, negative; orange, uncertain.

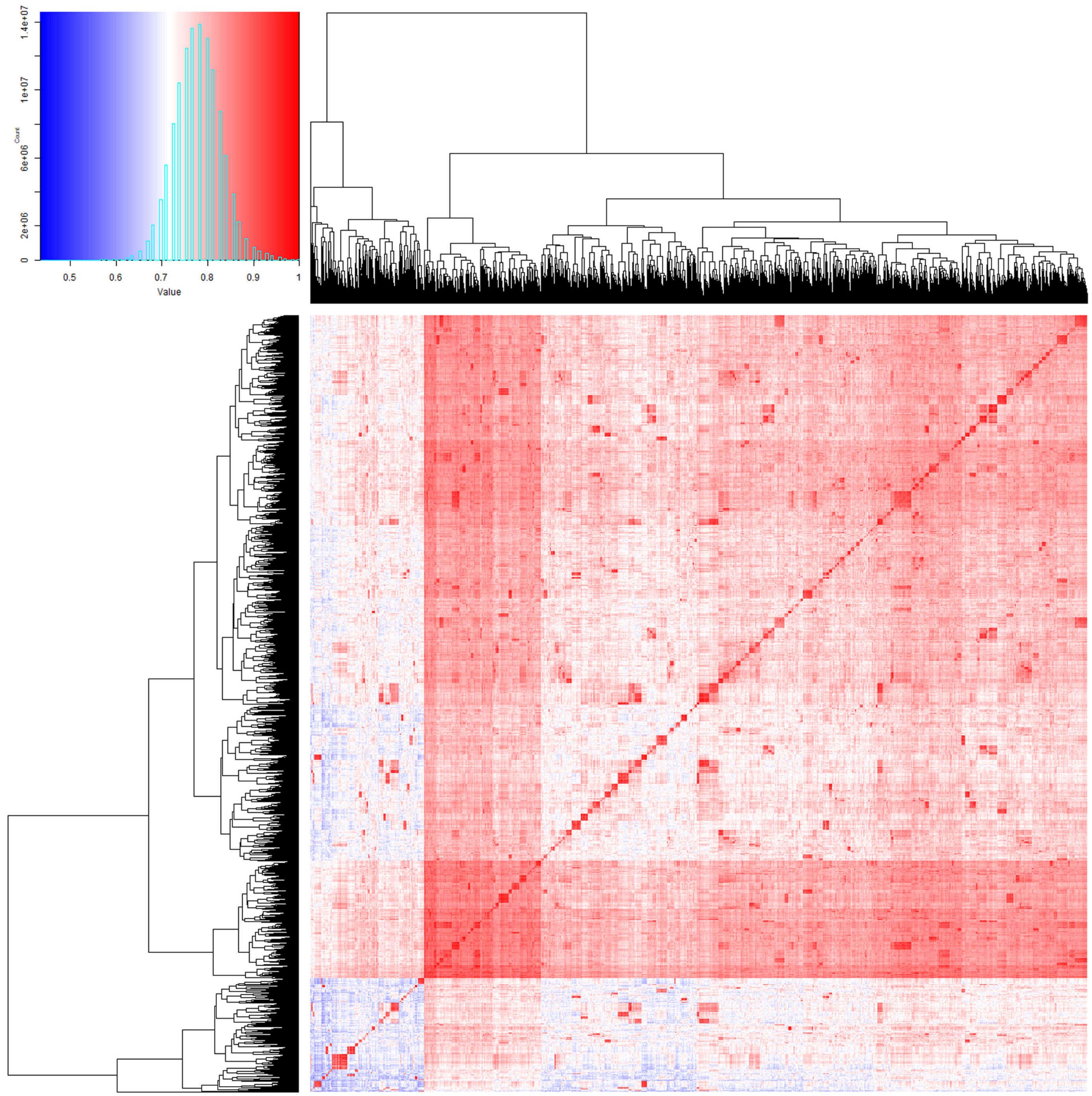


**Fig. S4.** Cluster analysis of beta satDNA sequences found in eukaryote genome assemblies. 10,966 full-length beta satDNA sequences were clustered and shown in heatmap using heatmap.2 in R language.

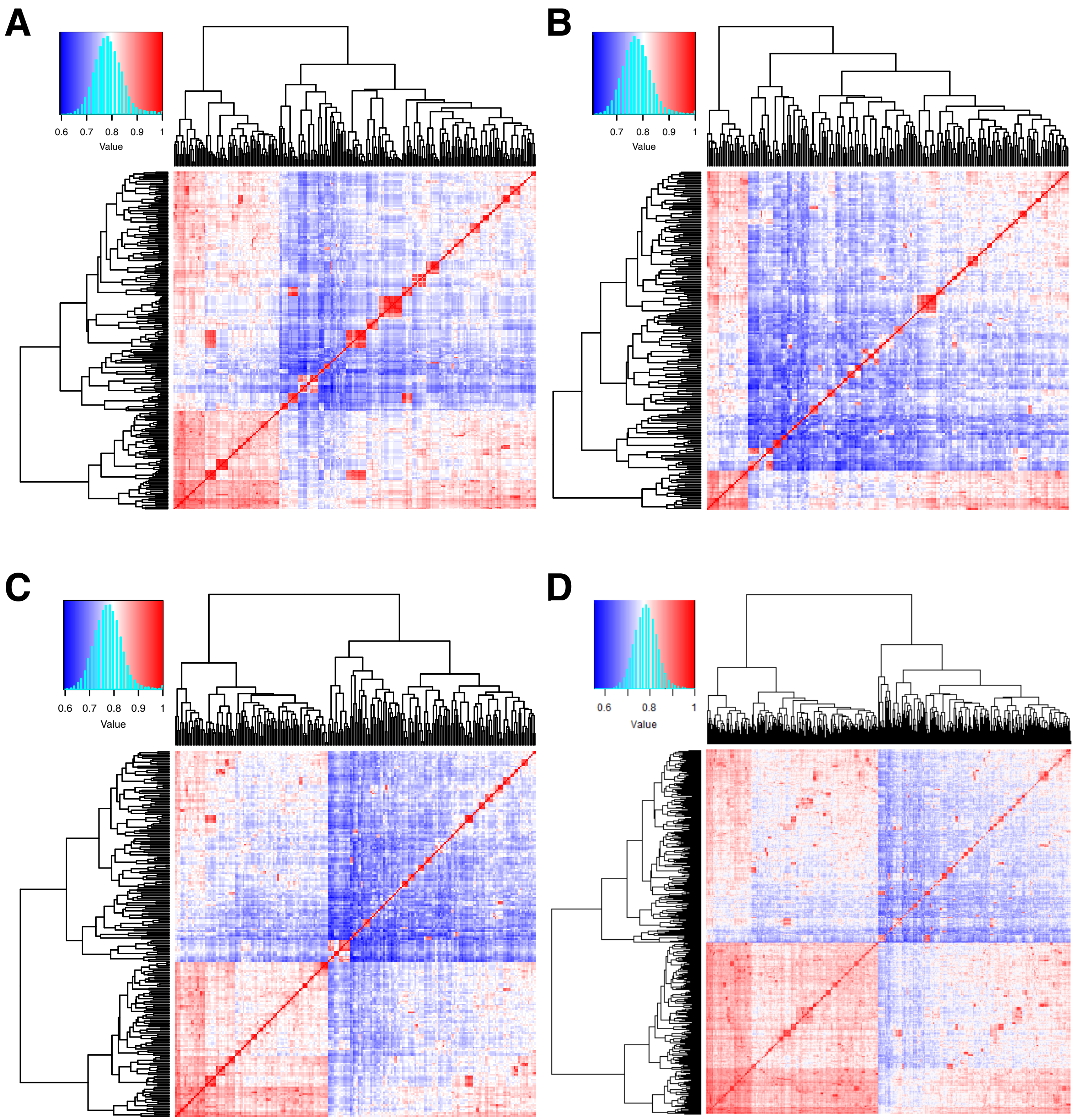


**Fig. S5.** Cluster analysis of beta satDNA sequences found in genome assemblies of different eukaryotic branches. A, animals excluding primates; B, fungus including mycetozoa; C, plants; D, Harosa. Sequences were clustered and shown in heatmap using heatmap.2 in R language.

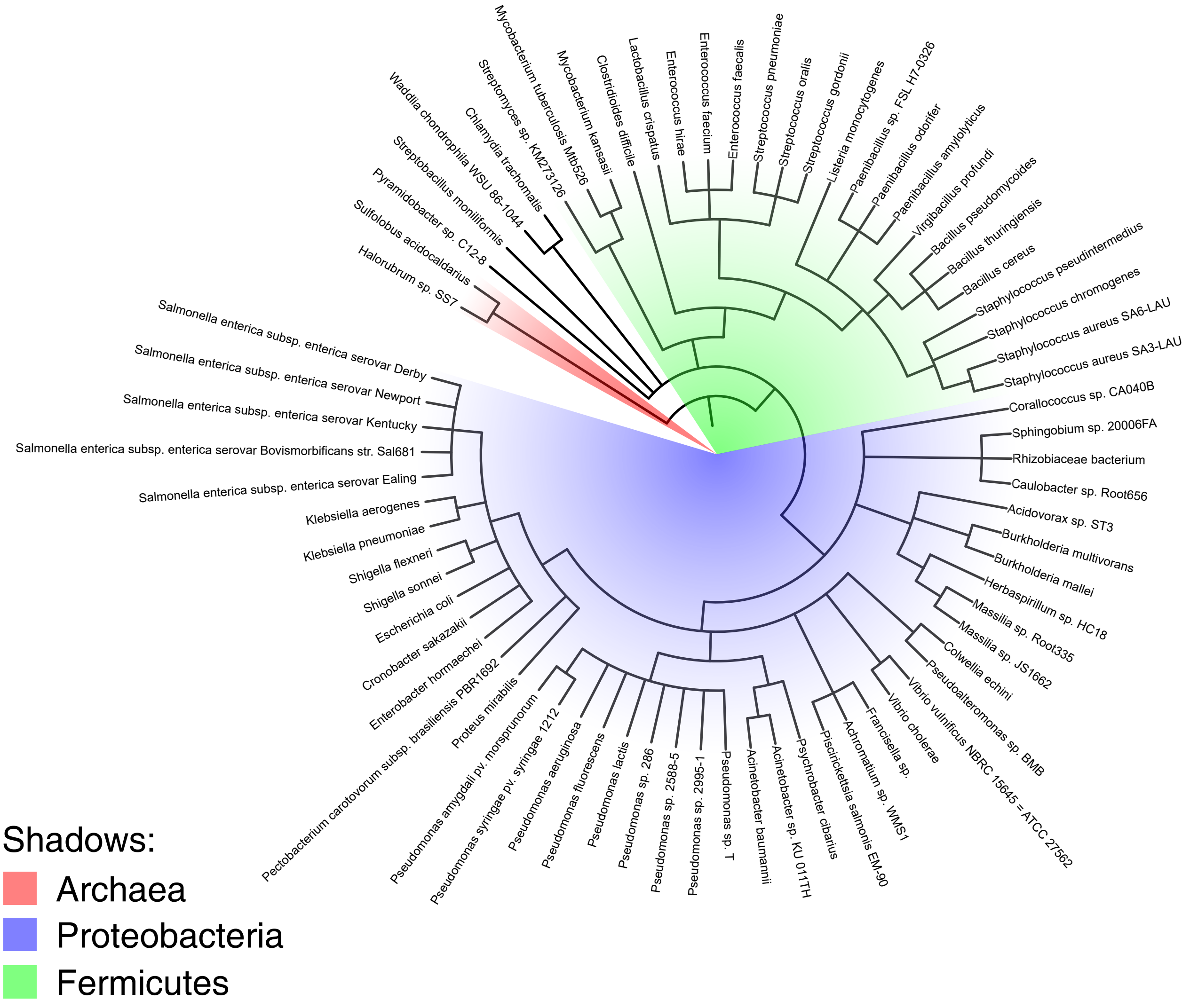


**Fig. S6.** Prokaryotic species containing beta satDNA sequences and their relationship. >210,000 prokaryotic genome assemblies were BLASTed against the beta satDNA index and 72 species were found containing beta satDNA sequences.

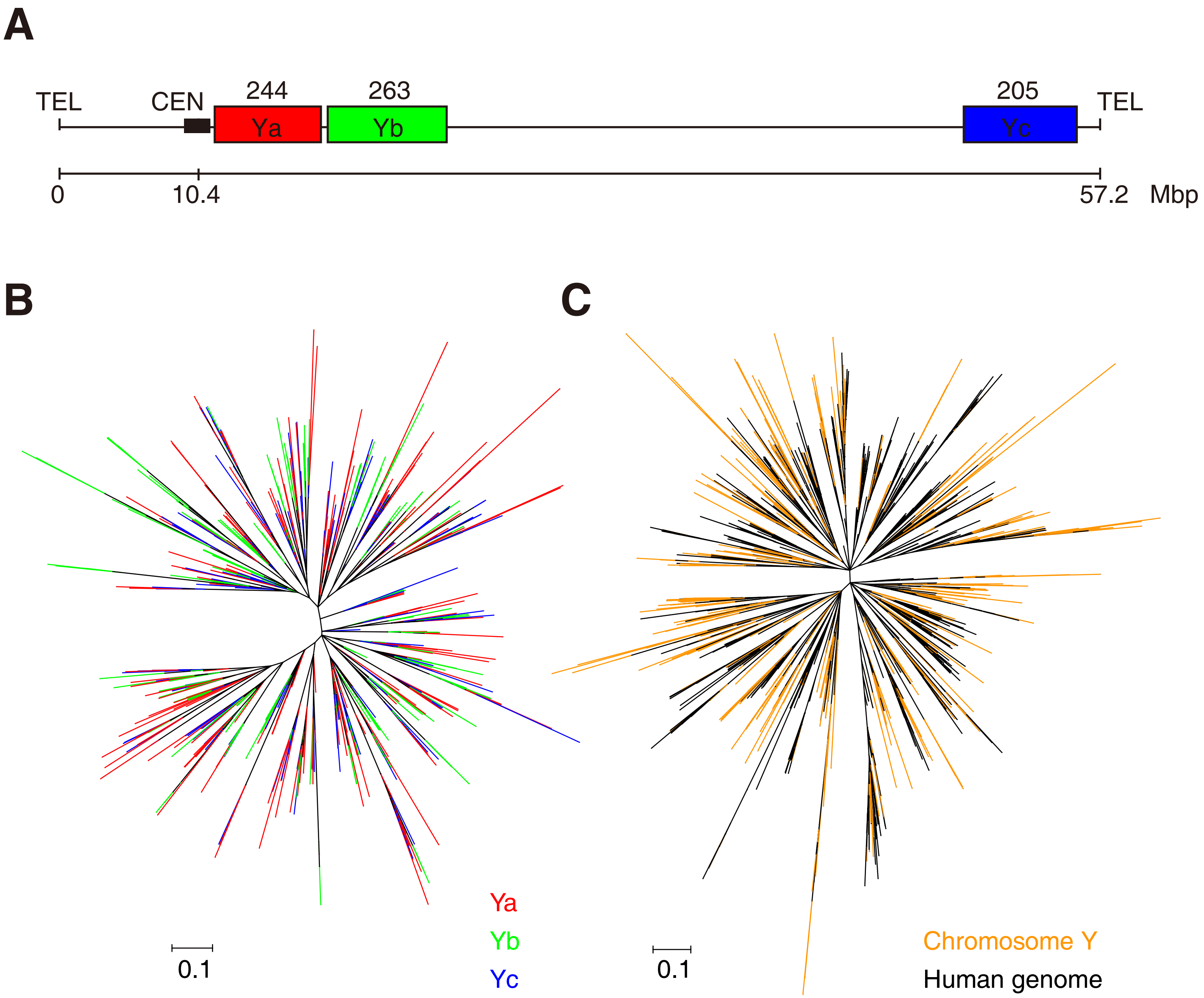


**Fig. S7.** Distribution and diversity of beta satDNA sequences on chromosome Y. A, Beta satDNA sequences on the chromosome Y. The beta satDNA copy numbers were labeled above the boxes. B, Phylogenetic tree of beta satDNAs on chromosome Y. Ya, Yb and Yc refer to beta satDNAs located at regions shown in a. C, Phylogenetic tree of beta satDNAs on chromosome Y (orange) and 1,000 copies beta satDNAs randomly picked from the human genome. The trees were constructed using RAxML with statistical support provided by bootstrapping over 1,000 replicates.

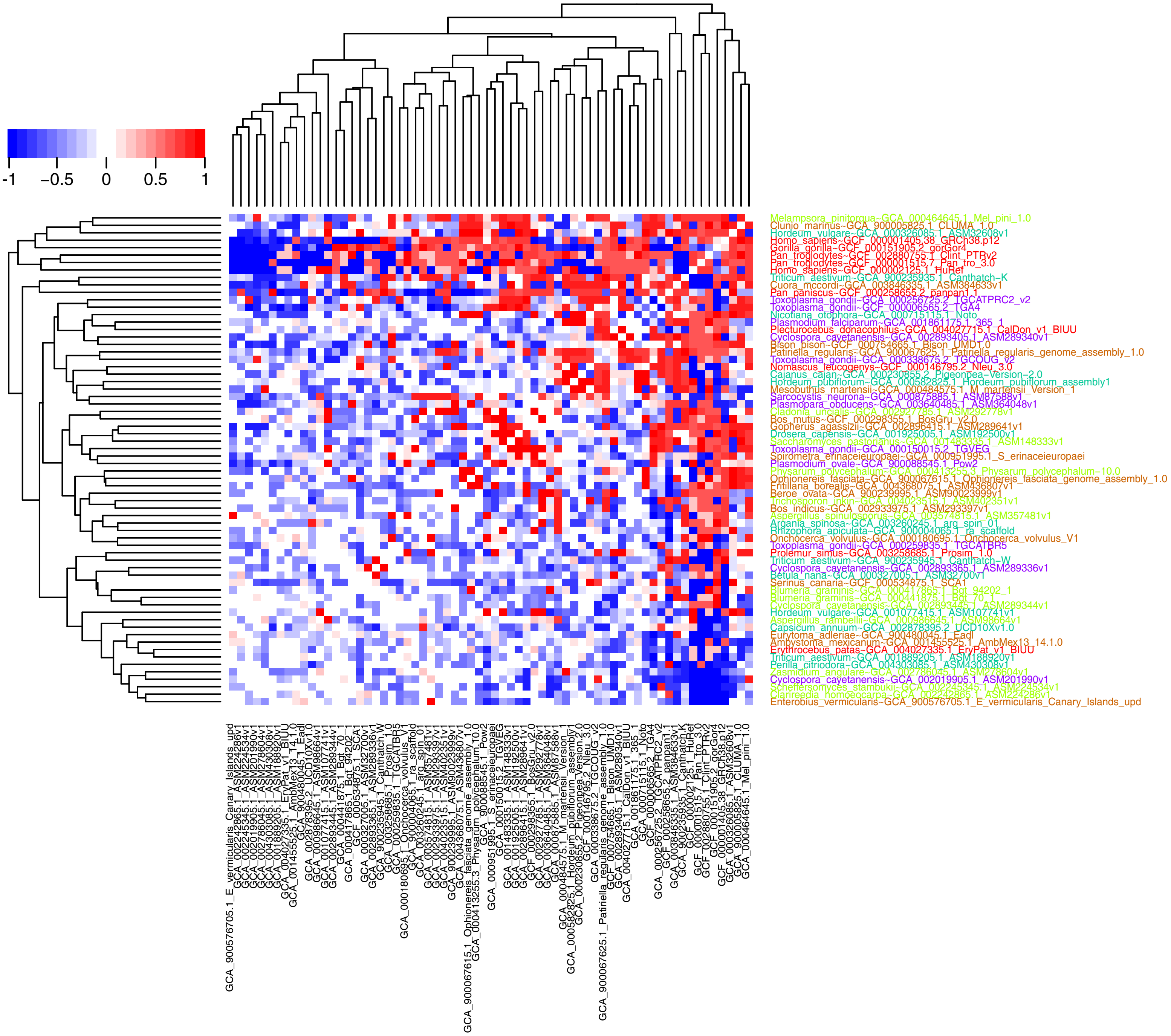


**Fig. S8.** Sequence clusters of libraries with randomized beta satDNA sequences. The sequences identified from the genome assemblies of different species (Supplementary File 1) were randomized and redistributed to the libraries again. Each library contains the same number of random sequences as the original library. Then the same analysis as Fig. 4B was performed and the heatmap was produced.

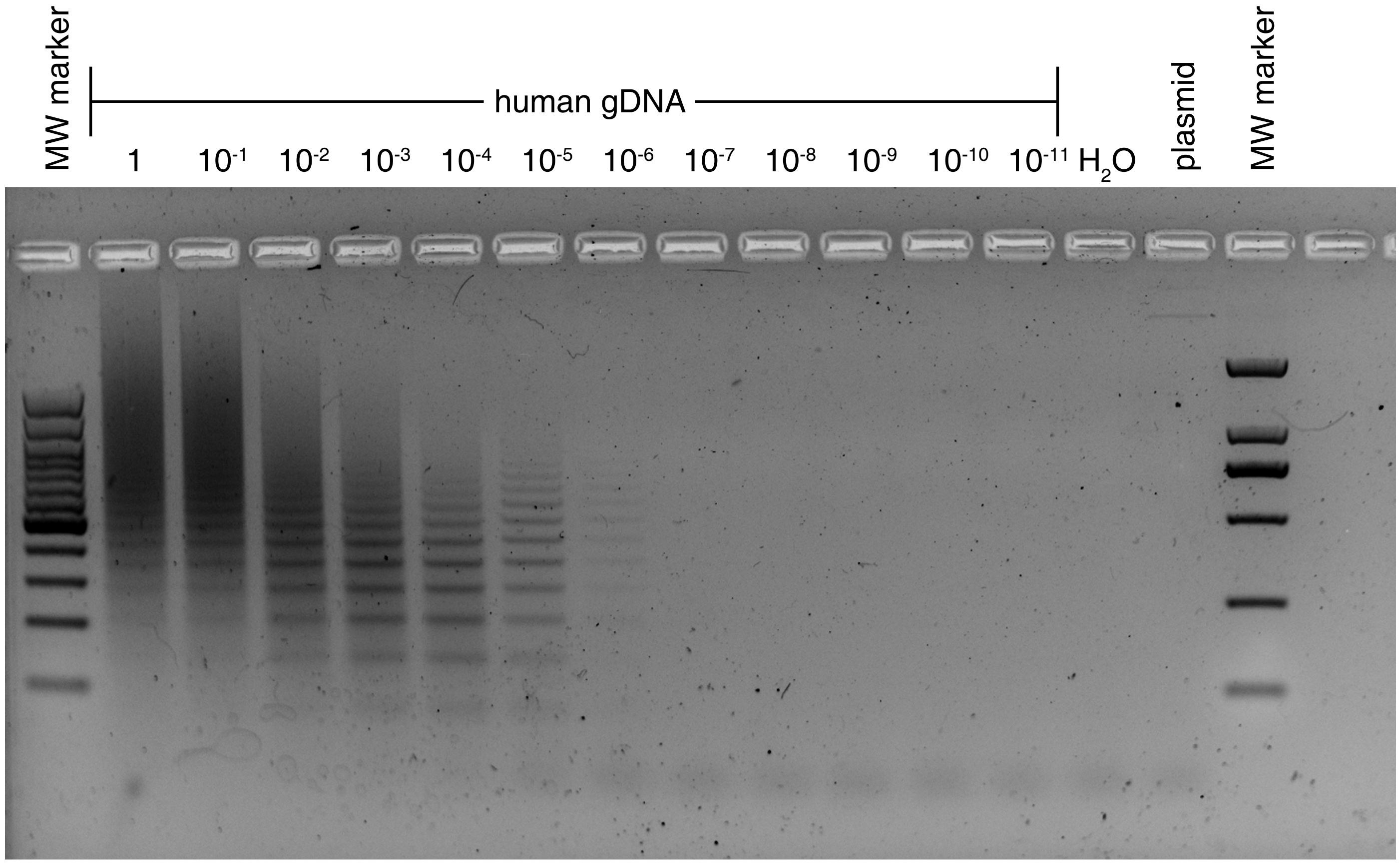


**Fig. S9.** PCR amplification using templates of consecutively diluted human gDNA. The original concentration of human genomic DNA was 5 ng/μl in the PCR system. Then serial dilutions were performed in 10-fold increments. The dilution range is from 1 to 1011 times. The PCR products were analyzed using electrophoresis with 1.5% agarose gel containing gel-red.

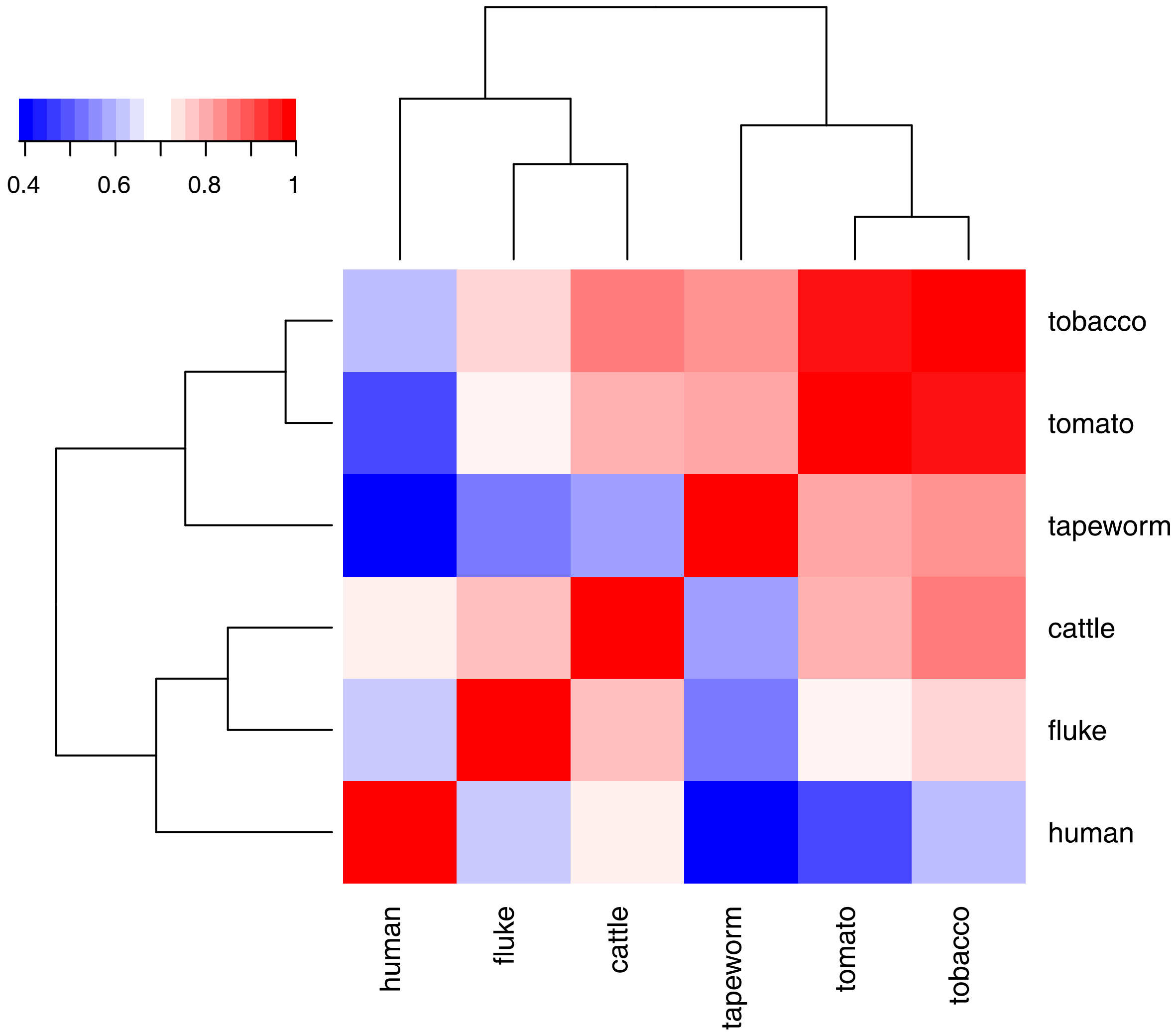


**Fig. S10.** Comparison of sequences obtained from next generation sequencing of PCR amplicons. The sequence reads of PCR amplicons were BLASTed against the beta satDNA index and the sequence libraries of different species were compared for common sequences. The matrix of common ratio was linearized and then used for generating heatmap in R language.
